# Abstract, modality-specific and experience-dependent coding of affect in the human brain

**DOI:** 10.1101/2023.08.25.554755

**Authors:** Giada Lettieri, Giacomo Handjaras, Elisa M. Cappello, Francesca Setti, Davide Bottari, Valentina Bruno, Matteo Diano, Andrea Leo, Carla Tinti, Francesca Garbarini, Pietro Pietrini, Emiliano Ricciardi, Luca Cecchetti

## Abstract

Emotion and perception are tightly intertwined, as affective experiences often arise from the appraisal of sensory information. Nonetheless, whether the brain encodes emotional instances using a sensory-specific code or in a more abstract manner is unclear. Here, we answer this question by measuring the association between emotion ratings collected during a unisensory or multisensory presentation of a full-length movie and brain activity recorded in typically-developed, congenitally blind and congenitally deaf participants. Emotional instances are encoded in a vast network encompassing sensory, prefrontal, and temporal cortices. Within this network, the ventromedial prefrontal cortex stores a categorical representation of emotion independent of modality and experience, and the posterior superior temporal cortex maps valence using an abstract code. Sensory experience more than modality impacts how the brain organizes emotional information outside supramodal regions, suggesting the existence of a scaffold for the representation of emotional states where sensory inputs during development shape its functioning.

## Introduction

The ability to comprehend and respond to affectively laden stimuli is vital and observing the behavior of others enables us to predict their reactions and tailor appropriate responses (Kunda, 1999; Frith and Frith, 2007). Interestingly, vision is our primary sensory modality to function in the external world, as it allows us to efficiently process a wealth of information from our surroundings (e.g., Eimer, 2004). Indeed, we heavily rely on this sense to interpret nonverbal visual cues coming from other individuals, such as facial expressions and body postures, which convey crucial information about an individual’s affective state (Itier and Batty, 2009; Tracy et al., 2015). Recent research in artificial intelligence and neuroimaging has highlighted the pivotal role of vision in understanding emotions, with convolutional neural networks predicting the emotional content of images, and the early visual cortex classifying distinct emotion categories (Kragel et al., 2019). In line with this, the same mechanism responsible for coding distinct stimulus properties in primary sensory areas (i.e., topographic mapping) supports the organization of affect in the right temporoparietal cortex (Lettieri et al., 2019).

While vision plays a dominant role in affective perception, we live in an environment that constantly engages multiple senses. Spoken communication, for instance, not only conveys semantic information but also provides cues about the speaker’s emotional state or intentions (Rieffe and Terwogt, 2000; Mozziconacci, 2002), as in the case of vocal bursts (Klinge et al., 2010; Cowen et al., 2019; Grollero et al., 2022). Furthermore, different emotional states exhibit unique auditory characteristics, with specific acoustic parameters correlating with arousal and valence (Banse and Scherer, 1996; Laukka et al., 2005; Bänziger et al., 2015).

Understanding the interplay between different sensory modalities in the representation and expression of emotions has been a subject of interest in behavioral, functional magnetic resonance imaging (fMRI), and electroencephalography studies (for a review see Schirmer and Adolphs, 2017). Specifically, multimodal presentations of emotional stimuli have been shown to enhance recognition accuracy and speed (Collignon et al., 2008; Klasen et al., 2012). This advantage may also reflect the brain organization, since previous studies have demonstrated that regions like the superior temporal sulcus and the prefrontal cortex successfully categorize emotions across different modalities (McCabe et al., 2008; Peelen et al., 2010; Chikazoe et al., 2014; Kim et al., 2015). Despite significant advances in the study of the interplay between perception and emotion, the majority of research have focused on static and unimodal emotional stimuli in typically-developed individuals only (e.g., Posner et al., 2009; Baucom et al., 2012). In this context, it is crucial to study congenital sensory deprivation, as this helps reduce the possibility that the activation of the same brain area across modalities can be solely attributed to mental imagery (Pietrini et al., 2004). In fact, in people with typical development, a fearful scream is likely to give rise to the mental imagery of someone in the act of screaming, and they are able to depict in their mind a specific facial expression and body posture, perhaps even a certain context, with a rich and dynamic representation close to what is commonly experienced in daily life. This process of mental imagery is associated with a specific pattern of brain activity in relation to different sensory channels (see for instance Farah, 1989; O’Craven and Kanwisher, 2000), and it can evoke vivid representations of emotional expressions and contextual information (Holmes and Mathews, 2010; Faul et al., 2022). Therefore, by using unimodal stimuli in typically-developed individuals, it becomes challenging to discern whether and where in the human brain emotional instances are represented using an abstract, rather than a sensory-dependent, code.

To address this question, congenital sensory deprivation constitutes a unique model, as individuals lacking information from a particular sensory channel, rely less on mental imagery associated with that channel (for a review, see Cattaneo et al., 2008). In this regard, previous evidence has shown that the brain’s representation of animacy (Pietrini et al., 2004; Gobbini et al., 2011; Ratan Murty et al., 2020), spatial layout (Wolbers et al., 2011), and objects (He et al., 2013) remains similar between typically-developed and sensory-deprived individuals, suggesting a supramodal encoding of information from the external world. However, it remains unclear whether this sensory-independent principle extends to the representation of affective states in the brain.

To address this gap, we conducted a study investigating the role of sensory experience and the contribution of mental imagery in the processing of emotional states. We collected moment-by-moment categorical and dimensional ratings of emotion in typically-developed individuals during an auditory-only, visual-only, or multisensory version of a full-length movie. We measured brain activity evoked by the same stimulus using fMRI in sighted and hearing participants and in those who were blind or deaf since birth. Our results reveal an abstract representation of categorical emotional states in the ventromedial prefrontal cortex and of the valence dimension in the posterior portion of the superior temporal gyrus. Additionally, we demonstrate that sensory experience more than modality impacts how the brain organizes emotional information outside supramodal regions, suggesting the existence of a scaffold for the representation of emotional states where sensory inputs during development shape its functioning.

## Materials and Methods

### Behavioral experiments - Participants

After having obtained their written informed consent, a sample of 124 Italian native speakers with typical visual and auditory experience (E^+^) participated in a set of behavioral experiments. All participants retained the right to withdraw from the study at any time and received a small monetary compensation for their participation. They had no history of neurological or psychiatric conditions, normal hearing, normal or corrected vision, and no reported history of drug or alcohol abuse.

A first group of 62 participants provided categorical ratings of the affective experience associated with the movie 101 Dalmatians under three experimental conditions: listening to the auditory-only version of the movie (auditory-only condition, A; n = 20, 8F, mean age ± standard deviation = 36 ± 16), watching the silent film (visual-only condition, V; n = 20, 10F, age = 29 ± 8), or watching and listening to the original version of the movie (multisensory condition, M; n = 22, 11F, age = 30 ± 3). A second independent sample of 62 individuals provided valence ratings of the emotional experience evoked by the same movie under the same experimental conditions. Specifically, 21 individuals reported the (un)pleasantness of the experience during the M condition (13F, age = 30 ± 5), 20 annotated the emotional valence during the A condition (9F, age = 37 ± 11), and 21 participated in the V experiment (10F, age = 29 ± 7). The mean age and the proportion of male and female individuals in each group did not differ from those of typically-developed and sensory deprived participants enrolled in the functional magnetic resonance imaging experiment (all p-values > 0.05). All participants had never watched the movie or had not watched it in the year prior to the experiment. The study was approved by the local Ethical Review Board (CEAVNO: Comitato Etico Area Vasta Nord Ovest; protocol no. 1485/2017) and conducted in accordance with the Declaration of Helsinki.

### Behavioral experiments - Stimuli and Experimental Paradigm

A shortened version of the 101 Dalmatians live-action movie (Walt Disney, 1996) was created for M, A, and V stimulation (Setti et al., 2023). Scenes irrelevant to the main plot were removed to limit the overall duration of the experimental session to one hour, and the movie was split into six runs for fMRI protocol compliance.

Six-second fade-in and fade-out periods were added at the beginning and the end of each run (iMovie software v10.1.10). A professional actor provided a voiceover of the story with a uniform pitch and no inflections in the voice, recorded in a studio insulated from environmental noise with professional hardware (Neumann U87ai microphone, Universal Audio LA 610 mk2 preamplifier, Apogee Rosetta converter, Apple MacOS) and software (Logic Pro). The voice track was then adequately combined with the original soundtracks and dialogues. Further, we added to each video frame subtitles of movie dialogues, text embedded in the video stream (e.g., newspaper), onomatopoeic sounds, and audio descriptions. Lastly, A and V versions of the movie were generated by discarding the video and audio streams, respectively.

Participants sat comfortably in a quiet room facing a 24" Dell™ screen, they wore headphones (Marshall™ Major III; 20–20,000 Hz; Maximum SPL 97 dB) and were presented with either the multimodal or unimodal edited versions of the movie 101 Dalmatians. Volunteers were asked to report their moment-to-moment emotional experience (10Hz sampling rate) using a collection of 15 emotion labels that were balanced between positive (amusement, joy, pleasure, contentment, love, admiration, relief, and compassion) and negative affective states (sadness, disappointment, fear, disgust, contempt, hate, and anger) states. As in our previous studies (Lettieri et al., 2019; 2022), the emotion labels were displayed on the bottom of the screen, evenly spaced along the horizontal axis, with positive emotion categories on the left and negative ones on the right side. Participants could navigate through the emotions using the arrow keys on a QWERTY keyboard, and once selected, the label changed its color from white to red. The onset or end of an emotional instance was marked by pressing the keys “Q” or “A”, respectively. Subjects were able to select multiple emotions at a time and could constantly monitor their affective reports based on the changing color of the corresponding label. For each individual, we obtained a 32,280 (i.e., timepoints) by 15 (i.e., emotion labels) matrix of affective ratings, in which a value of 1 (or 0) indicated the presence (or absence) of a specific emotion at a given time. Prior to the experiment, participants received training on the task, in which a label appeared on the screen (e.g., joy), and they were asked to reach the corresponding emotion category and mark the onset as fast as they could. They were instructed to proceed to the actual experiment only if they felt comfortable performing the task.

We followed a similar procedure to collect real-time ratings of valence (i.e., dimensional ratings). This time, however, participants were provided with a single “valence” label and were asked to evaluate how pleasant or unpleasant their experience was at each moment by increasing (“Q” keypress) or decreasing (“A” keypress) the valence score. The valence scale ranged from -100 (extremely unpleasant) to +100 (extremely pleasant), the minimum step was set to 5 points, and a value of 0 indicated a neutral state. Participants were able to monitor their affective state in real-time and adjust the valence score at any moment (10Hz sampling frequency). We collected a timeseries of 32,280 timepoints for each individual, with positive or negative values indicating the pleasantness or unpleasantness of their emotional experience at any given time.

Both behavioral experiments were conducted using MATLAB (R2019b; MathWorks Inc., Natick, MA, USA) and Psychtoolbox v3.0.16 (Kleiner et al., 2007).

### Behavioral experiments - Data Analysis

Single-participant matrices of categorical affective ratings were downsampled to the fMRI temporal resolution (i.e., 2 seconds) and then aggregated across individuals by summing the number of occurrences of each emotion at each timepoint. The resulting 1,614 (i.e., downsampled timepoints) by 15 (i.e., emotions) matrix stored values ranging from 0 to the maximum number of participants experiencing the same emotion at the same time. To account for idiosyncrasies in group-level affective ratings, timepoints in which only one participant reported experiencing a specific emotion were set to 0. The group-level matrix of categorical ratings was then normalized by dividing values stored at each timepoint by the overall maximum agreement in the matrix. Because these ratings were used to explain the brain activity of independent participants in a voxelwise encoding analysis, we convolved the 15 group-level emotion timeseries with a canonical hemodynamic response function (spm_hrf function; HRF) and added the intercept to the model. The entire procedure was repeated for ratings collected in the M, the A, and the V conditions. Similarly, single-participant valence ratings were downsampled to the fMRI temporal resolution and aggregated at the group level by computing the average valence score across individuals at each timepoint and convolved with a canonical HRF. These processing steps were applied to valence ratings obtained from the M, the A, and the V experiments.

To evaluate the between-participant similarity in categorical ratings of emotion, for each volunteer and condition, we obtained the emotion-by-timepoint matrix expressing the onset and duration of emotional instances throughout the movie. We then computed the Jaccard similarity between all pairings of individuals. This index quantified the proportion of timepoints in which two volunteers reported the experience of the same state, over the number of timepoints in which either individuals felt the emotion (0 = completely disjoint reports, 1 = perfect correspondence). The overlap between emotional reports was then summarized by computing the median across all possible pairings of participants in each condition and emotion category. For valence ratings, instead, the between-participant similarity was evaluated using Spearman’s *ρ* and summarized by computing the median of correlation values obtained from all pairings of participants in each condition. Representational similarity analysis (RSA; Kriegeskorte et al., 2008) was used to measure whether group-level emotion ratings collected during movie watching recapitulated the arrangement of emotion categories within the space of affective dimensions. In this regard, we first obtained from a large database of affective norms (Warriner et al., 2013) the scores of valence, arousal, and dominance dimensions (1 to 9 Likert scale; Russell and Mehrabian, 1977) of each emotion label. A representational dissimilarity matrix (RDM) was obtained by computing the between-emotion pairwise Euclidean distance in the space of affective dimensions. We then used Kendall’s *τ* to correlate the affective norms (RDM) with behavioral RDMs obtained from emotion ratings of each experimental condition (i.e., M RDM, A RDM, and V RDM). Behavioral RDMs were derived by estimating the Spearman’s correlation *ρ* between all possible pairings of group-level emotion timeseries. In addition to evaluating the association between the affective norms RDM and each behavioral RDM, we also measured Kendall’s *τ* correlation between all pairings of behavioral RDMs, hence testing the correspondence between affective ratings collected under diverse sensory modalities.

To further prove the specificity of categorical ratings across conditions, we tested whether the distance (Spearman’s *ρ* on group-level ratings) of a specific emotion acquired under two different experimental conditions (e.g., amusement M versus amusement A) was smaller than its distance to all other emotions (e.g., amusement M versus any other emotion A). This produced a non-symmetric emotion-by-emotion RDM summarizing the distance between emotion ratings acquired under two modalities and was repeated for all condition pairings (i.e., M versus A, M versus V, and A versus V). For each emotion and RDM, we then identified the category with the minimum distance, thus allowing a direct inspection of the consistency of emotion ratings between sensory modalities.

Lastly, to reveal the structure of categorical ratings and test their correspondence with valence scores collected in behavioral experiments, we performed principal component (PC) analysis on the group-level emotion-by-timepoints matrix. The number of components was set to three to comply with the affective norms by Warriner and colleagues (2013), and the procedure was repeated on data obtained from the M, A, and V experiments. Of each component, we inspected the variance explained and the coefficients of emotion categories. The timecourse of each component (i.e., PC scores) was also extracted and correlated (Kendall’s *τ*) with the behavioral valence scores collected under the same experimental condition (e.g., PC1 M versus valence M). In addition, we also reported Kendall’s *τ* correlation between all possible condition pairings in terms of valence ratings and scores of the principal components. The strength of Spearman’s and Kendall’s correlations was interpreted following the recommendation from Schober and colleagues (2018).

### fMRI experiment - Participants

Brain activity evoked by the same movie employed in the behavioral experiments was measured in a group of 50 Italian volunteers. Participants were categorized into five groups based on their sensory experience and on the stimulus presentation modality: a group of congenitally blind individuals listening to the auditory-only version of 101 Dalmatians (E^‒^A; n = 11, 3F, age 46 ± 14), a sample of congenitally deaf without cochlear implants watching the silent film (E^‒^V; n = 9, 5F, age 24 ± 4), and three groups of typically-developed individuals, who were presented with either the auditory-only (E^+^A; n = 10, 7F, 39 ± 17), the visual-only (E^+^V; n = 10, 5F, 37 ± 15), or the multisensory (E^+^M; n = 10, 8F, 35 ± 13) versions of the stimulus. During the fMRI acquisition, participants were instructed to remain still and enjoy the movie. The scanning session lasted approximately 1 hour and was divided into six functional runs (3T Philips Ingenia scanner, Neuroimaging center of NIT - Molinette Hospital, Turin; 32 channels head coil; gradient recall echo-echo planar imaging - GRE-EPI; 2000ms repetition time, 30ms echo time, 75° flip angle, 3 mm isotropic voxel, 240 mm field of view, 38 sequential ascending axial slices, 1614 volumes). Audio and visual stimulations were delivered using MR-compatible LCD goggles and headphones (VisualStim Resonance Technology, video resolution 800x600 at 60 Hz, visual field 30° × 22°, 5′′, audio 30 dB noise-attenuation, 40 Hz to 40 kHz frequency response). High-resolution anatomical images were also acquired (3D T1w; magnetization-prepared rapid gradient echo; 7ms repetition time, 3.2ms echo time, 9° flip angle, 1mm isotropic voxel, 224 mm field of view). The fMRI study was approved by the Ethical Committee of the University of Turin (Protocol No. 195874/2019), and all participants provided their written consent for participation.

### fMRI experiment - Preprocessing

For each participant, the high-resolution T1w image was brain extracted (OASIS template, antsBrainExtraction.sh) and corrected for inhomogeneity bias (N4 bias correction) with ANTs v2.1.0 (Avants et al., 2011a). The anatomical image was then non-linearly transformed to match the MNI152 ICBM 2009c non-linear symmetric template using AFNI v17.1.12 (3dQwarp; Cox, 1996). Also, binary masks of white matter (WM), grey matter (GM), and cerebrospinal fluid (CSF) were obtained from Atropos (Avants et al., 2011b). We used these masks to extract the average timeseries of WM and CSF voxels from fMRI sequences, which were included in the functional preprocessing pipeline as regressors of no interest (Ciric et al., 2017). Masks were transformed into the MNI152 space by applying the already computed deformation field (3dNwarpApply; interpolation: nearest neighbor). To ensure that after normalization to the standard space the WM mask included WM voxels only, we skeletonized the mask (erosion: 3 voxels; 3dmask_tool) and excluded (3dcalc) the Harvard-Oxford subcortical structures (i.e., thalamus, caudate nucleus, pallidum, putamen, accumbens, amygdala, and hippocampus). Similarly, the CSF mask was eroded by 1 voxel and multiplied by a ventricle mask (MNI152_T1_2mm_VentricleMask.nii.gz) distributed with the FSL suite (Smith et al., 2004). Lastly, both masks were downsampled to match the fMRI spatial resolution. For each participant and functional run, we corrected slice-dependent delays (3dTshift) and removed non-brain tissue (FSL bet -F) from the images. Head motion was compensated by aligning each volume of the functional run to a central timepoint (i.e., TR: 134) with rigid-body transformations (i.e., 6 degrees of freedom; 3dvolreg). The motion parameters, the aggregated timeseries of absolute and relative displacement, and the transformation matrices were generated and inspected. Also, we created a brain-extracted motion-corrected version of the functional images by estimating the average intensity of each voxel in time (3dTstat). This image was coregistered to the brain-extracted anatomical sequence (align_epi_anat.py, “giant move” option, lpc+ZZ cost function). To transform the functional data into the MNI152 space using a single interpolation step, we concatenated the deformation field, the coregistration matrix, and the 3dvolreg transformation matrices and applied the resulting warp (3dNwarpApply) to the brain-extracted functional images corrected for slice-dependent delays. Standard-space functional images were generated using sinc interpolation (5 voxels window), having the same spatial resolution as the original fMRI data (i.e., 3mm isotropic voxel). Brain masks obtained from functional data were transformed into the standard space as well (3dNwarpApply, nearest neighbor interpolation). Functional images were then iteratively smoothed (3dBlurToFWH) until a 6mm full width at half maximum level was reached. As in Ciric and colleagues (2017), we employed 3dDespike to replace outlier timepoints in each voxel’s timeseries with interpolated values and then normalized the signal so that changes in blood oxygen level-dependent (BOLD) activity were expressed as a percentage (3dcalc). In addition, WM, CSF, and brain conjunction masks were created by identifying voxels common to all functional (3dmask_tool -frac 1). We selected 36 predictors of no interest (36p) to be regressed out from brain activity, which included 6 head motion parameters (6p) obtained from 3dvolreg, average timeseries (3dmaskave) of WM (7p), average CSF signal (8p), and the average global signal (9p). For each of these nine regressors, we computed their quadratic expansions (18p; 1deval), the temporal derivatives (27p; 1d_tool.py), and the squares of derivatives (36p). Also, regressors of no interest were detrended using polynomial fitting (up to the 5^th^ degree). Confounds were regressed out using a generalized least squares timeseries fit with restricted maximum likelihood estimation of the temporal autocorrelation structure (3dREMLfit). Regression residuals represented single-participant preprocessed timeseries, which were then employed in voxelwise encoding, univariate comparisons, multivariate classification and cross-decoding analyses.

### fMRI experiment - Voxelwise encoding

We utilized a voxelwise encoding approach to identify brain regions that are involved in the representation of affect across different sensory modalities and in individuals with varying sensory experiences. In this regard, categorical emotion ratings from the A, V, and M versions of 101 Dalmatians served as predictors of brain activity recorded in independent participants watching and/or listening to the same stimulus. Specifically, A ratings were used as the encoding model of fMRI data collected in congenitally blind (i.e., E^‒^A) and typically-developed individuals (i.e., E^+^A) listening to the stimulus; emotions timeseries obtained from the V experiment were fitted to the brain activity of congenitally deaf (i.e., E^‒^V) and typically-developed participants (i.e., E^+^V) watching the silent version of the movie; lastly, affective ratings coming from the M behavioral experiment served to explain changes in the BOLD activity of E^+^M volunteers attending the multisensory version of the stimulus.

The encoding analysis was performed at the single-participant level, thus producing a voxelwise R^2^ map for each typically-developed and sensory-deprived individual. The statistical significance of the full-model fit was established using a non-parametric permutation approach. In brief, the rows of the encoding matrix were shuffled 2,000 times prior to the HRF convolution. Therefore, the resulting null encoding matrices preserved the co-occurrence of emotion categories found in actual ratings, whereas the onset and duration of emotional instances were randomized. The null encoding matrices were then convolved with a canonical HRF and fitted to brain activity to produce 2,000 null R^2^ maps for each participant. Importantly, the same permutation scheme was used for all participants and voxels. In each voxel, the position of the unpermuted R^2^ value in the null distribution determined the voxelwise uncorrected statistical significance level (i.e., p-value). Because we were interested in group-level encoding maps of affect, we then used a non-parametric combination (Winkler et al., 2016; Fisher method) to aggregate p-values across participants of the same group. We opted for this approach because, in the case of a one-sample test and unsigned statistics (e.g., R^2^, F-stat), the traditional sign-flip method produces unreliable estimates of statistical significance. Instead, the non-parametric combination measures convergence of significance across participants, thus revealing brain regions encoding changes in affect in each group and modality. The group-level significance was then adjusted for the number of comparisons using the cluster-based method (cluster-determining threshold: p-valueCDT < 0.001 uncorrected) and the family-wise correction (FWC) suggested by Nichols and Holmes (2002; p-valueFWC < 0.05).

### fMRI experiment - Univariate contrasts and conjunction analyses

To study if sensory experience and stimulus modality exert impact on the mapping of affective states in the brain, we have conducted four univariate unpaired t-tests on voxelwise encoding R^2^ values and four conjunction analyses on binary maps of significance. Two univariate tests compared the congenitally blind and deaf participants with their matching groups of typically-developed individuals listening to or watching the movie (i.e., equation a: E^‒^A ≠ E^+^A; equation b: E^‒^V ≠ E^+^V). A third test (i.e., equation c) was aimed at unveiling brain areas involved in the mapping of affect in both sensory-deprived groups, but not in typically-developed people, as a consequence of shared crossmodal reorganizations: (E^‒^A + E^‒^V) ≠ (E^+^A + E^+^V). Lastly, a fourth test (i.e., equation d) revealed the regions in which the representation of affect did not depend on the sensory experience but was specific to the sensory modality: (E^‒^V + E^+^V) ≠ (E^‒^A + E^+^A). For each voxel and comparison, first, the unpermuted average group difference in the fitting of the emotion model was estimated using a pseudo-t statistic. Then, we randomly permuted 2,000 times participants’ identities and computed the pseudo-t under the null hypothesis of no group differences. The uncorrected p-values were determined by the position of the unpermuted pseudo-t in the null distribution, and a cluster-based correction was applied to account for the number of comparisons (p-value_CDT_ < 0.001, p-value_FWC_ < 0.05; Nichols and Holmes, 2002). As far as conjunction analyses are concerned, the first measured the intersection between the five binary maps of group-level significance obtained from the voxelwise encoding procedure (i.e., equation e: E^+^M ∩ E^+^A ∩ E^+^V ∩ E^‒^A ∩ E^‒^V). This conjunction revealed brain areas (if any) mapping affect regardless of the sensory modality and experience, namely a supramodal (Cecchetti et al., 2016) representation of emotion. A second and third conjunction analyses assessed the overlap between the typically-developed and the sensory-deprived groups within modality (i.e., equation f: E^+^A ∩ E^‒^A; equation g: E^+^V ∩ E^‒^V). Lastly, a fourth conjunction determined the spatial correspondence between voxels encoding affect across modalities in typically-developed people (i.e., equation h: E^+^A ∩ E^+^V).

### fMRI experiment - Multivoxel pattern classification analysis

We conducted a multivoxel classification analysis to test whether brain regions represent emotion with a distinctive pattern that is informed by the stimulus modality and/or by the participant’s sensory experience. Firstly, we selected all brain areas significantly encoding the emotion model in at least two groups. This brain mask determined the voxels entered as features in a 5-class linear support vector machine (SVM; class labels: E^+^M, E^+^A, E^+^V, E^‒^A, E^‒^V; soft margin parameter C = 1) aimed at classifying sensory experience and stimulus modality from the single-participant R^2^ encoding maps. The classification was performed using a 5-fold cross-validation procedure, and the features were standardized across participants (i.e., z-score transformation). At each iteration, participants’ data were split into training (n = 40) and test (n = 10) sets, and a minimum redundancy maximum relevance (MRMR) algorithm was used to select the most informative 1,000 features (i.e., voxels) in the training set, thus reducing the risk of overfitting. Also, the reason for implementing the feature selection step at each iteration of the k-fold cross-validation was to measure the contribution of each voxel to the classification (i.e., the number of times each voxel was considered informative across the folds). The model parameters of the multiclass SVM classifier were estimated in the training set and then applied to the test set to predict the class identity of the left-out observations. The same procedure was repeated for all the folds, and a confusion matrix was built to report the matching between actual and predicted class labels. The global performance of the multiclass SVM was assessed using the weighted F1-score, which takes into account both precision and recall and is robust to class imbalance. The statistical significance of the classification was assessed by repeating the entire procedure (i.e., cross-validation, standardization, feature selection, estimation of model parameters in the training set, and prediction in the test set) on data for which the participants’ sensory experience and stimulus modality (i.e., class labels) were randomized 2,000 times. We obtained p-values relative to the global performance of the classifier as well as to every single class.

### fMRI experiment - Cross-decoding analysis

To characterize the information content of brain regions involved in the representation of emotion, we used a cross-decoding approach and predicted ratings of hedonic valence from the brain activity of sensory-deprived and typically-developed participants. In brief, we selected brain areas significantly encoding the emotion model in at least two groups (as in the multivoxel pattern analysis) and used their activity in a L2 penalized regression (i.e., ridge regression; optimization method: stochastic gradient descent) to explain average valence ratings obtained from independent participants. The association between brain activity and valence was first estimated with fMRI and behavioral data acquired under the same condition (i.e., stimulus modality) and in people with a specific sensory experience and then tested in all other groups and conditions. Ridge regression coefficients were determined using a leave-one-out cross-validation procedure. Specifically, we left out the fMRI data of one participant (e.g., an E^‒^A observation; validation set) and averaged data collected under the same condition and from all other participants with the same sensory experience (e.g., all other E^‒^A observations; training set). Ridge regression coefficients were estimated for penalization factors in the range 1*10^-3^ < λ < 1*10^2^ (1,000 logarithmically spaced values) and then applied to the left-out observation. The mean squared prediction error (MSE) of valence scores was obtained for each penalization factor and left-out fMRI participant, and the optimal λ was established by minimizing the average MSE. Ridge regression coefficients relative to the optimal cross-validated penalization factor were then obtained in the complete training group (e.g., all E^‒^A observations) and applied to all other groups (i.e., test sets) to cross-decode valence ratings. For instance, the optimal ridge coefficients for the prediction of auditory-only valence scores from the brain activity of congenitally blind individuals were obtained and then used to predict multisensory valence scores from the brain data of typically-developed individuals listening to and watching the original version of the movie. The prediction was tested at the single-participant level, and the statistical significance was assessed through timepoint shuffling of valence ratings (n = 2,000 iterations), which led to permutation-based estimates of the prediction error (i.e., MSE) under the null hypothesis. Single-participant results were then aggregated at the group level using the non-parametric combination approach (Winkler et al., 2016; Fisher method). The entire procedure was repeated for all groups and conditions.

## Results

### Behavioral experiments

Firstly, we explore the distribution of group-level categorical reports of emotion based on the 15 labels. Amusement, love, and joy are the positive emotions more frequently used to describe the affective experience across the multisensory (M), the auditory-only (A) and the visual-only (V) modalities. Negative states, instead, are more often labeled as fear or contempt (Figure 1a-b, e-f, and i-j). Using PC analysis, we show that the first component, which contrasts positive and negative states, explains most of the variance in categorical ratings: 36.0% in the M condition, 41.2% in the A condition, and 43.5% in the V condition (Figure 1c, g, and k).

**Figure 1.**
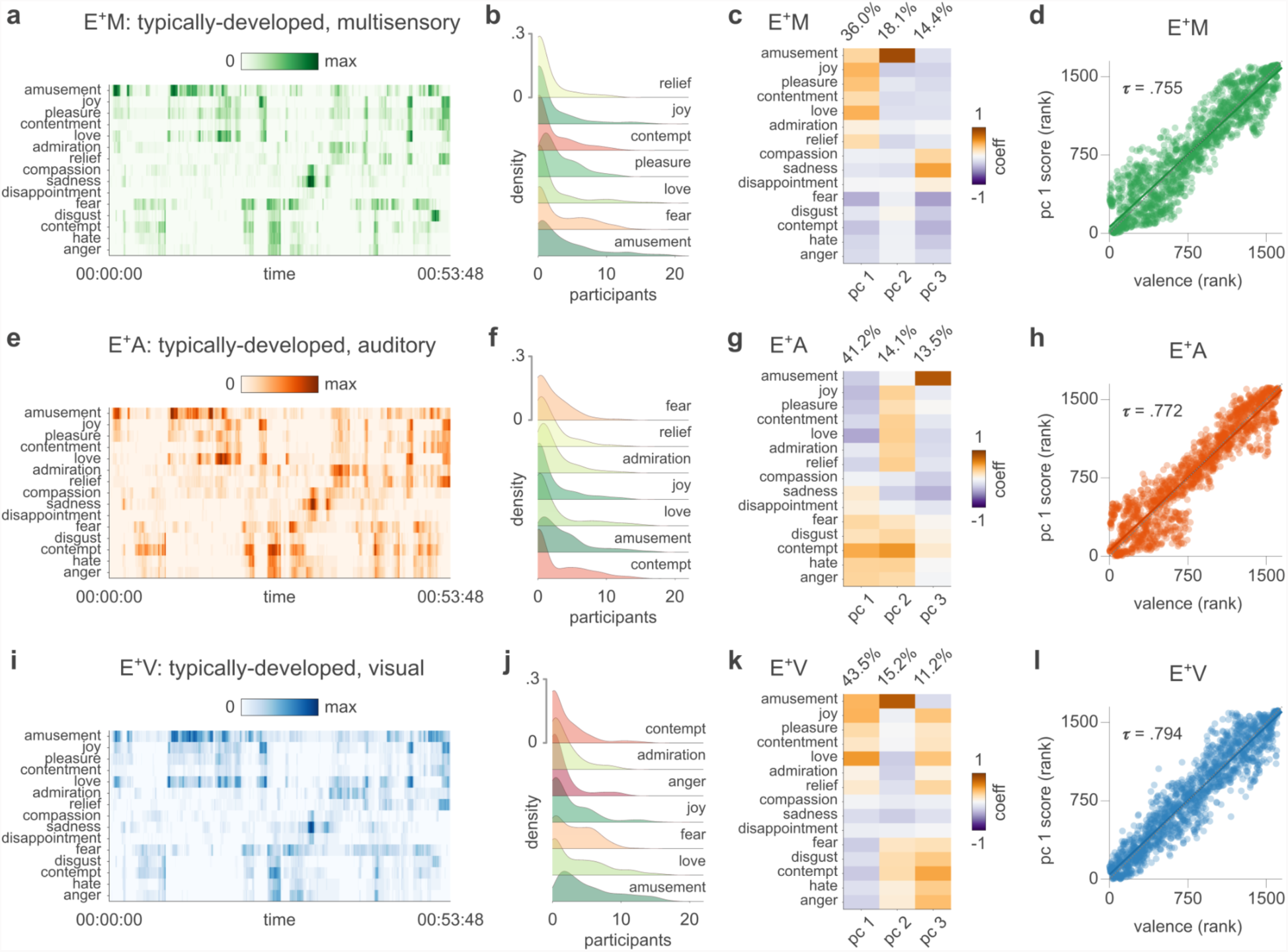
Emotion ratings across sensory modalities. Panel **a** shows the group-level emotion ratings collected from typically-developed participants in the multisensory condition (E^+^M). Darker colors indicate that a higher proportion of volunteers have reported the same emotion at a given point in time. Panel **b** depicts the distribution of movie timepoints (i.e., density) as a function of the between-participant overlap in categorical ratings for the first 7 emotions. In **c**, we show the loadings and the explained variance of the first three principal components obtained from the emotion-by-timepoint group-level matrix. In **d**, we report the correlation between the first principal component and valence ratings collected in independent participants. Panels **e-h** and **i-l** summarize the same information for the auditory (E^+^A) and the visual (E^+^V) conditions, respectively.

Also, in each modality, average valence ratings obtained from independent participants very strongly correlate with the scores of the first PC extracted from categorical reports (M valence ∼ M PC 1: *τ* = 0.755, M valence ∼ M PC 2: *τ* = 0.138, and M valence ∼ M PC 3: *τ* = 0.012; A valence ∼ A PC 1: *τ* = 0.772, A valence ∼ A PC 2: *τ* = 0.046, and A valence ∼ A PC 3: *τ* = 0.038; V valence ∼ V PC 1: *τ* = 0.794, V valence ∼ V PC 2: *τ* = 0.018, and V valence ∼ V PC 3: *τ* = 0.034; Figure 1d, h, and l). The analysis of the between-participant agreement in categorical ratings reveals that the median Jaccard similarity index is *J* = 0.245 for sadness, *J* = 0.225 for amusement, *J* = 0.181 for love, and *J* = 0.158 for joy in the M condition. In the A condition, the highest median between-participant correspondence in categorical ratings is observed for amusement (*J* = 0.183), followed by contempt (*J* = 0.177), love (*J* = 0.176), and sadness (*J* = 0.167). Lastly, in the V condition, amusement (*J* = 0.229), sadness (*J* = 0.172), joy (*J* = 0.152), and love (*J* = 0.147) are the categories used more similarly by participants. As far as the valence ratings are concerned, median between-participant Spearman’s correlation is moderate for all conditions (M: *ρ* = 0.483, A: *ρ* = 0.572, V: *ρ* = 0.456). Results of the RSA show that group-level emotion ratings collected under the three experimental conditions relate to the arrangement of emotion labels within the space of affective norms (Figure 2a). Specifically, the pairwise distance between emotions in the valence-arousal-dominance space correlates moderately with the RDM built from categorical ratings collected in the M condition (*τ* = 0.419) and strongly with those relative to the A (*τ* = 0.556) and the V (*τ* = 0.485) modalities. Moreover, there is a very strong correlation between RDMs obtained from the three modalities, particularly between the two unimodal conditions (M ∼ A: *τ* = 0.734, M ∼ V: *τ* = 0.777, A ∼ V: *τ* = 0.812).

**Figure 2.**
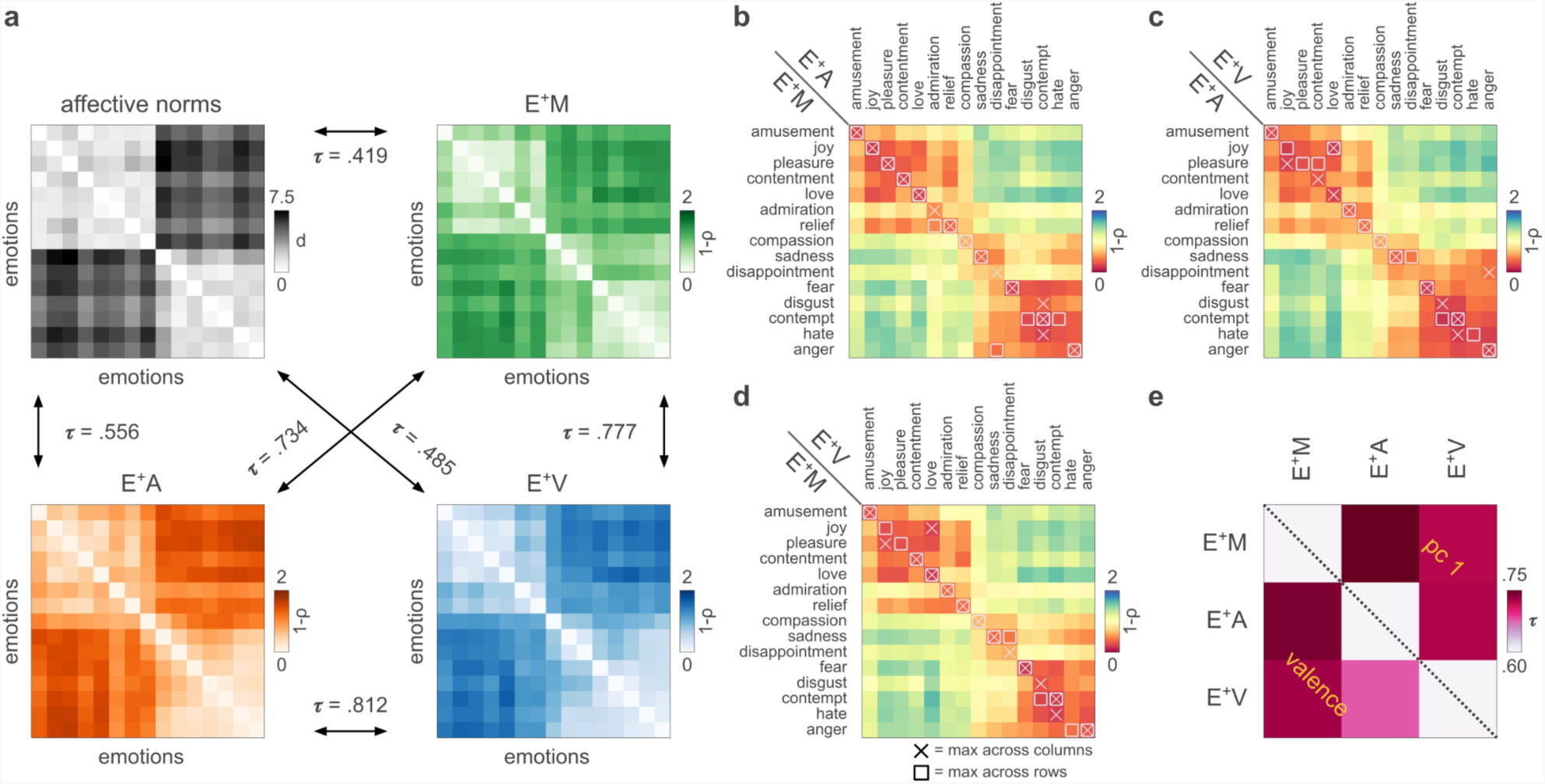
Similarity in emotion ratings between modalities and their relation with affective norms. Panel **a** shows that group-level emotion ratings collected under the three experimental conditions correlate with each other and relate to the arrangement of emotions in the space of affective norms. In **b**, **c**, and **d**, we show that, in most of the cases, ratings of a specific emotion acquired under one condition correlate maximally with ratings of the same emotion acquired under a different condition. The lower triangular part of the matrix in panel **e** depicts the correlation in valence between modalities, while the upper triangular part shows the correlation in terms of PC1 scores derived from categorical reports.

Further exploring the specificity of categorical ratings of emotion across sensory modalities, we demonstrate that, in the vast majority of cases, the correlation between the timecourse of a specific emotion acquired under two distinct conditions is higher than its correlation with all other emotions (Figure 2b-d). Importantly, when pairs of emotions are confounded between modalities they are also similar in valence (e.g., joy and love) and semantically related (e.g., contempt, hate, and disgust).

Lastly, measuring the correlation between valence ratings collected under different sensory modalities (Figure 2e; lower triangular part), we observe strong associations for the M vs A (*τ* = 0.742) and the M vs V (*τ* = 0.728) conditions, as well as a moderate correlation between the two unimodal conditions (A ∼ V: *τ* = 0.679). When this analysis is repeated on principal component scores from categorical ratings (Figure 2e; upper triangular part), we confirm the strength of the M vs A (*τ* = 0.766) and the M vs V (*τ* = 0.721) relationships, and reveal a strong correlation between the two unimodal conditions as well (A ∼ V: *τ* = 0.723).

### fMRI experiment - Voxelwise encoding

By examining voxelwise encoding results in typically-developed individuals presented with the multisensory version of the movie (i.e., E^+^M), we observe that the emotion category model is encoded bilaterally in the lateral orbitofrontal cortex (lOFC), the ventromedial prefrontal cortex (vmPFC), the medial prefrontal cortex (mPFC), and the anterior portion of the bilateral superior temporal gyrus (aSTG). Also, the affective experience is mapped in the central and posterior segments of the right superior temporal gyrus and sulcus (STG/STS), the right superior parietal lobule (SPL), and the right inferior occipital gyrus (IOG; p-value_FWC_ < 0.05, Figure 3a).

**Figure 3.**
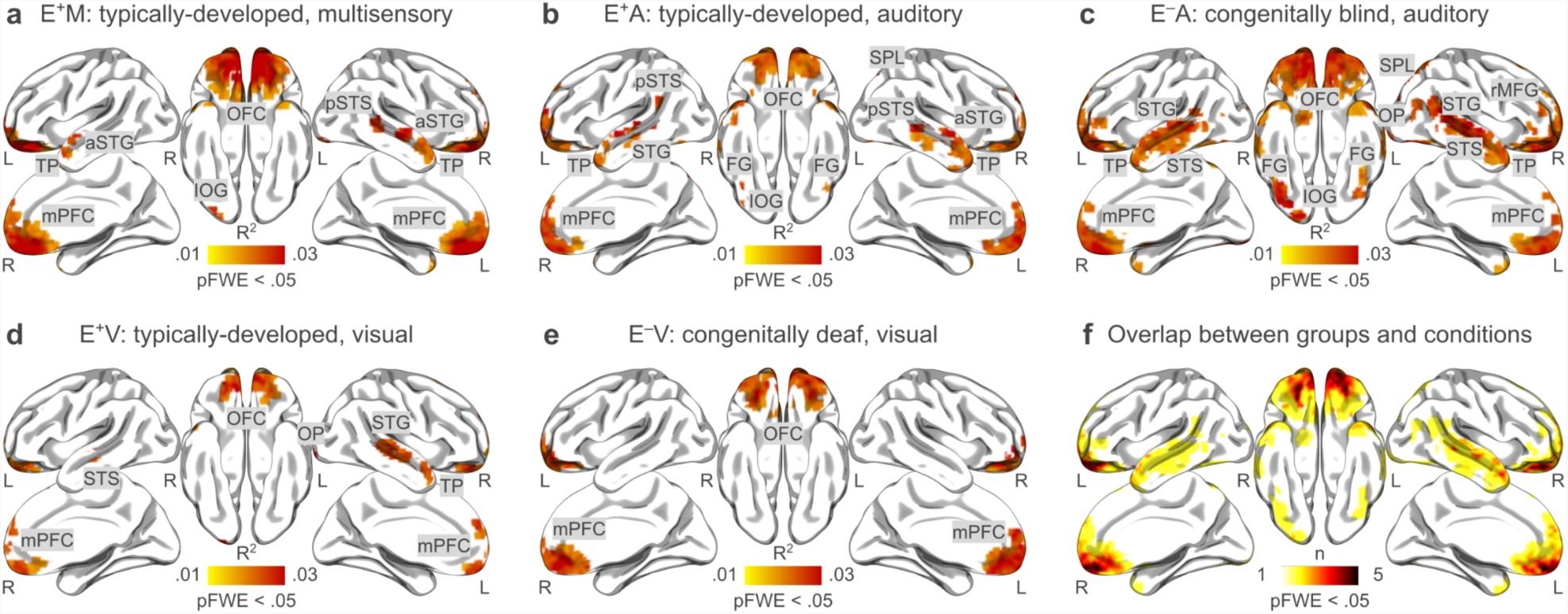
Group-level voxelwise encoding of the emotion model. Panels **a-e** show the results of the group-level voxelwise encoding analysis after correction for multiple comparisons (cluster-based correction, cluster forming threshold: p < .001; family-wise threshold: p < .05). Ratings of 15 distinct emotion categories provided by 62 E^+^ (M: n = 22; A: n = 20; V: n = 20) watching and/or listening to the 101 Dalmatians live action movie were used to explain fMRI activity recorded in E^+^ (panel **a**, M: n = 10; panel **b**, A: n = 10; panel **d**, V: n = 10) and E^‒^ (panel **c**, A: n = 11; panel **e**, V: n = 9) people presented with the same movie. Panel **f** shows the overlap of the group-level encoding results (pFWE < .05) between all groups and conditions. aSTG = anterior superior temporal gyrus, FG = fusiform gyrus, IOG = inferior occipital gyrus, mPFC = medial prefrontal cortex, OFC = orbitofrontal cortex, OP = occipital pole, pSTS = posterior superior temporal sulcus, rMFG = rostral middle frontal gyrus, SPL = superior parietal lobule, STG = superior temporal gyrus, STS = superior temporal sulcus, TP = temporal pole, L = left hemisphere, R = right hemisphere.

When typically-developed individuals listen to the audio version of the movie (i.e., E^+^A), emotion categories are represented bilaterally in the lOFC, the mPFC, the vmPFC, the STG/STS, the IOG, the SPL, the fusiform gyrus (FG), and in the left supramarginal gyrus (SMG; p-value_FWC_ < 0.05, Figure 3b). Similarly, the brain of congenitally blind individuals (i.e., E^‒^A) represents emotions in the bilateral mPFC, the vmPFC, the lOFC, the STG/STS, the IOG, and the FG. In addition to these regions, the activity of the right lingual gyrus (LG), the right occipital pole (OP), the right ventral diencephalon (vDC), and the bilateral rostral middle frontal gyrus (rMFG) encodes the affective experience in congenitally blind people listening to the movie (p-value_FWC_ < 0.05, Figure 3c).

In the video-only condition, emotional instances are mapped mainly in the bilateral lOFC, the mPFC, the vmPFC, the right STG/STS, the right LG, and the right OP of the typically-developed brain (E^+^V; p-value_FWC_ < 0.05, Figure 3d). Concerning the results obtained from congenitally deaf individuals (i.e., E^‒^V), the affective experience is represented in the bilateral lOFC, the mPFC, the vmPFC, and the left STG/STS (p-value_FWC_ < 0.05, Figure 3e). Overall, the mPFC, the vmPFC, the OFC, and the STS represent emotion categories across the majority of conditions and in people with varying sensory experience (Figure 3f).

### fMRI experiment - Univariate contrasts and conjunction analyses

Univariate comparisons between people with and without sensory deprivation - i.e., E^‒^A ≠ E^+^A; E^‒^V ≠ E^+^V; (E^‒^A + E^‒^V) ≠ (E^+^A + E^+^V) - show no significant differences in the extent to which individual voxels encode the emotion model (all clusters p-value_FWC_ > 0.05). Instead, we observe that, regardless of sensory experience, emotion categories are encoded in the bilateral auditory cortex with larger fitting values when people are presented with the audio-only version of the movie - i.e., (E^‒^V + E^+^V) < (E^‒^A + E^+^A) - (Figure 4a; Table 1). At the same time, the emotion model fits more the bilateral early visual cortex when typically-developed and congenitally deaf people watch the silent movie - i.e., (E^‒^V + E^+^V) > (E^‒^A + E^+^A) - (Figure 4a; Table 1).

**Figure 4.**
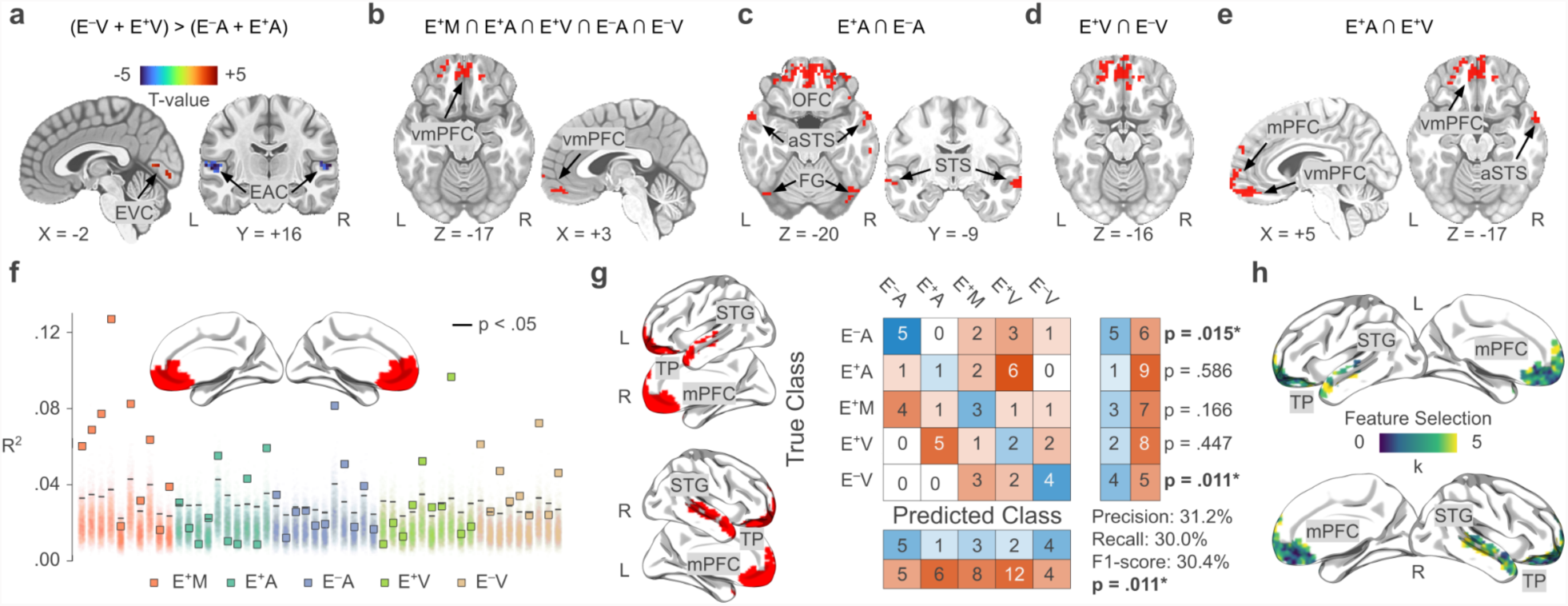
Univariate and multivariate analyses of the emotion network and its association with sensory modality and experience. Panel **a** depicts regions encoding the emotion model as a function of the sensory modality. In red, voxels showing higher fitting values in the visual modality, and in blue those more specific for the auditory one. Panels **b**-**e** summarize the results of conjunction analyses. Areas highlighted in red in panel **b** map the emotion model regardless of the sensory experience and modality. In **c** and **d**, we show the overlap between typically-developed and sensory-deprived individuals presented with the auditory and the visual stimulus, respectively. Panel **e** represents the convergence between voxels encoding affect in the two unisensory modalities in people with typical experience. Panel **f** shows single-participant results of the association (full model R^2^) between emotion ratings and average activity of vmPFC defined using Neurosynth. Squares represent the fitting of the emotion model in each participant (typically-developed multisensory - E^+^M: red; typically-developed auditory - E^+^A: cyan; congenitally blind auditory - E^‒^A: blue; typically-developed visual - E^+^V: green; congenitally deaf visual - E^‒^V: brown). Shaded areas refer to the single-participant null distributions and solid black lines represent the p<.05 significance level. In **g**, we show the results of the multivoxel pattern classification analysis. Voxelwise encoding R^2^ maps are used to predict the participant sensory experience and stimulus modality. The central part of the panel shows the confusion matrix and the performance of the multiclass (n=5; chance ∼20%) cross-validated (k=5) SVM classifier. Overall, classification performance is significantly different from chance (p=.011) and driven by the successful identification of sensory-deprived individuals. Feature importance analysis (panel **h**) shows that voxels of vmPFC were rarely (or never) selected to predict sensory experience and modality. OFC=orbitofrontal cortex, mPFC=medial prefrontal cortex, STG=superior temporal gyrus, STS=superior temporal sulcus, aSTS=anterior superior temporal sulcus, TP = temporal pole, FG=fusiform gyrus, vmPFC=ventromedial prefrontal cortex, EVC=early visual cortex, EAC=early auditory cortex, L=left hemisphere, R=right hemisphere.

**Table 1.**
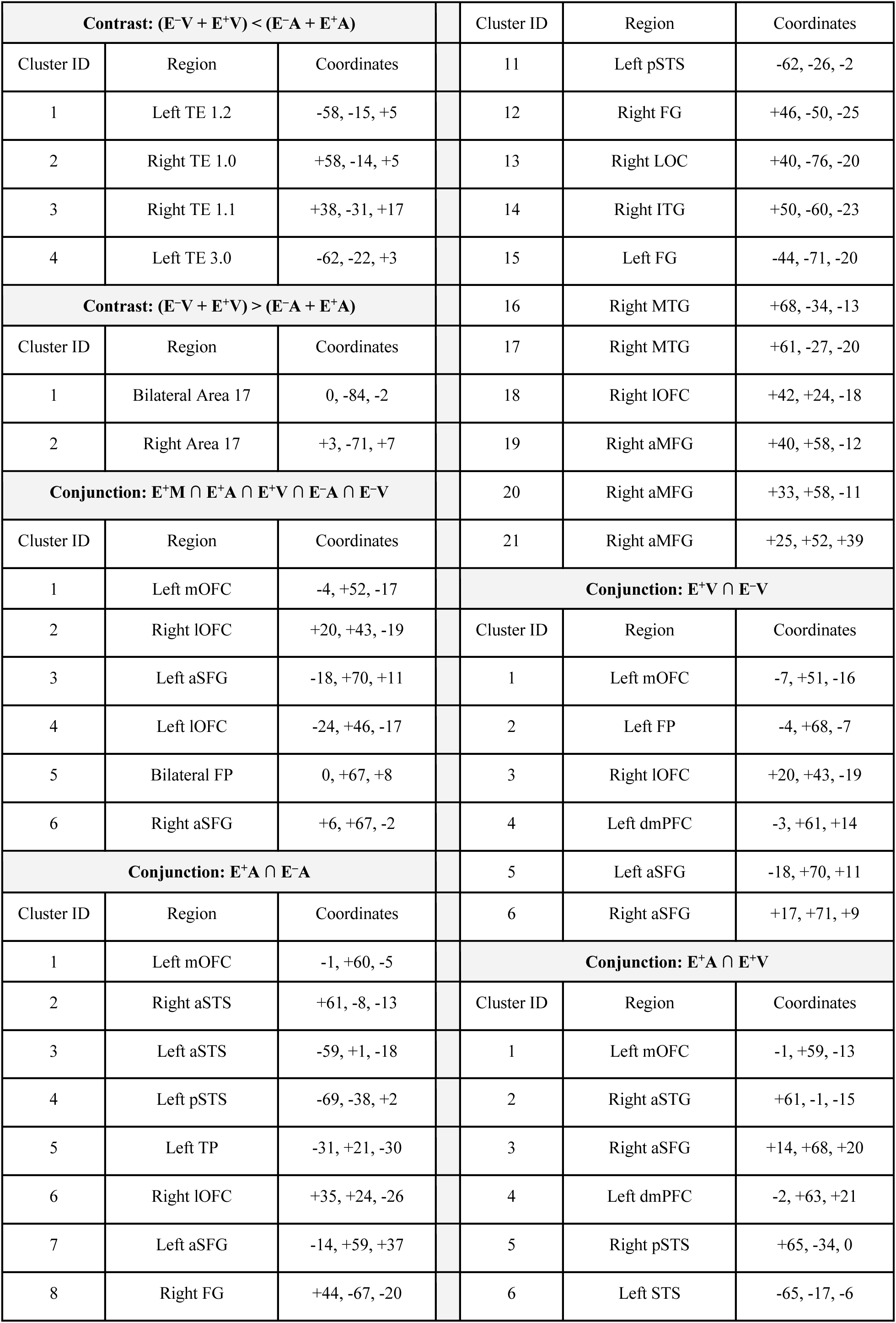
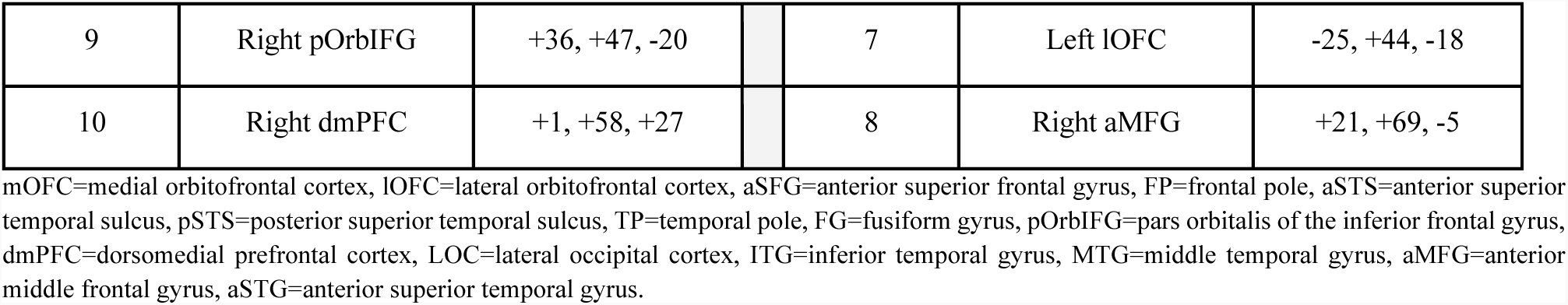
Results of univariate contrasts and conjunction analyses.

The conjunction analysis between groups and conditions demonstrates that the vmPFC, the FP, and the lOFC all map emotions regardless of the sensory experience and stimulus modality (Eq. e; Figure 4b; Table 1). Additionally, we show that the overlap between regions encoding affect in blind and sighted individuals listening to the audio-only movie extends to the bilateral temporal and occipital cortex, such as the STG/STS, and the FG (Eq. f; Figure 4c; Table 1). Instead, the conjunction between normally-hearing and congenitally deaf volunteers watching the silent movie reveals that only medial prefrontal areas are involved in the mapping of emotion categories (Eq. g; Figure 4d; Table 1). Lastly, when considering voxels encoding affect across modalities in typically-developed people, we observe convergence in frontal and temporal regions (Eq. h; Figure 4e; Table 1).

To further show that the activity of the vmPFC is associated with emotion ratings at the single-participant level and avoid double-dipping (Kriegeskorte et al., 2009), we obtain a mask of this region from the website version of Neurosynth (https://neurosynth.org/; Yarkoni et al., 2011; term: “vmpfc”, association test) and extract the average vmPFC signal across voxels in each participant. The resulting time-series is used as a dependent variable in a general linear model having the emotion ratings as predictors. The strength of the association between the categorical model of emotion and the vmPFC average activity is measured using the coefficient of determination R^2,^ and statistical significance is assessed by permuting the timepoints of the predictor matrix 2,000 times.

Results show that the emotion model significantly relates to the average vmPFC activity (p-value < 0.05) in 28 out of 50 participants (Figure 4f): 8 E^+^M, 5 E^+^A, 4 E^‒^A, 4 E^+^V, and 7 E^‒^V.

### fMRI experiment - Multivoxel pattern classification analysis

As an alternative to the univariate approach, we test in a multivariate fashion whether the pattern obtained from the fitting of the emotion model provides sufficient information to identify the sensory experience of participants and the stimulus modality they are presented with. Findings show above-chance classification (F1-score: 30.4%, Precision: 31.2%, Recall: 30.0%; p-value = 0.011, Figure 4g), particularly when it comes to identifying sensory-deprived individuals. Indeed, the 5-class classifier correctly recognizes 45.5% of E^‒^A individuals (p-value = 0.015) and 44.4% of E^‒^V volunteers (p-value = 0.011). Instead, we fail to predict the sensory modality to which typically-developed individuals are exposed to (E^+^A: 10.0%, p-value = 0.586; E^+^V: 20.0%, p-value = 0.447; E^+^M: 30.0%, p-value = 0.166). Also, the feature importance analysis reveals that, while voxels of bilateral dmPFC and left anterior STS contribute the most to the classification, those of bilateral vmPFC are never selected by the algorithm (Figure 4h).

### fMRI experiment - Cross-decoding analysis

To characterize the information content of brain regions significantly encoding the emotion model (Figure 5a), we estimate the association between hemodynamic activity and valence ratings in each condition and group and then test whether the relationship holds in all other conditions and groups. Results show that the cross-decoding of hedonic valence from the whole emotion network is possible for all pairings of conditions and groups, with the exception of E^+^A (Figure 5b). Specifically, using regression weights estimated in typically-developed individuals listening to the movie, we successfully explain the association between brain activity and valence in congenitally blind individuals (E^‒^A, p-value = 0.007), but not in other groups (E^+^M, p-value = 0.075; E^+^V, p-value = 0.738; E^‒^V, p-value = 0.820). In line with this, the relationship between the hemodynamic activity and valence scores in E^+^A can be reconstructed from regression coefficients estimated in blind people exclusively (E^‒^A, p-value = 0.001; E^+^M, p-value = 0.117; E^+^V, p-value = 0.180; E^‒^V, p-value = 0.358).

**Figure 5.**
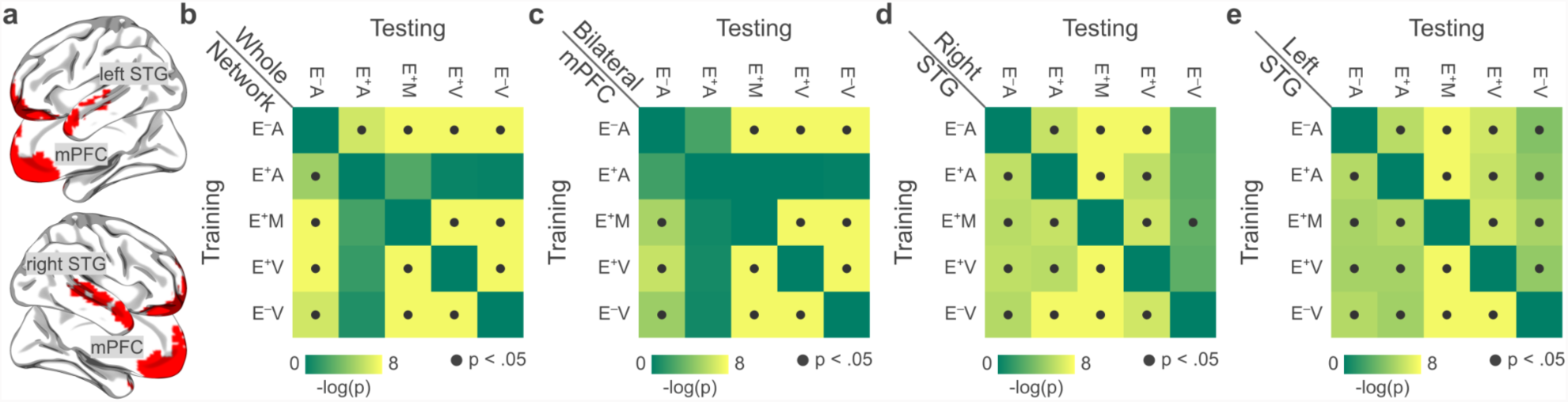
Cross-decoding of valence ratings from activity of regions encoding the emotion model. Panel **a** depicts the brain regions included in the cross-decoding procedure. In **b**-**e**, we show the results of the cross-validated ridge regression aimed at predicting valence ratings from brain activity of the emotion network (**b**), the bilateral mPFC (**c**), the right STG (**d**), and the left STG (**e**), acquired in all conditions and groups. We trained the algorithm (i.e., identified ridge coefficients) on each group and condition (training, matrix rows) and tested the association between brain activity and valence in all other groups and conditions (testing, matrix columns). Results are summarized by the 5-by-5 matrices showing the significance of the prediction for each pairing (dark gray dots denote p < .05). mPFC = medial prefrontal cortex, STG = superior temporal gyrus.

To further improve the spatial specificity of the cross-decoding results, we conduct the same analysis on each region of the emotion network separately. Similar to what we report for the entire network, apart from E^+^A, cross-decoding is possible for all pairings of conditions and groups in the bilateral mPFC cluster (Figure 5c). In this case, however, we have not been able to reconstruct the relationship between brain activity and valence in congenitally blind people starting from regression weights of typically-developed individuals listening to the movie (p-value = 0.131), nor vice versa (p-value = 0.105). As far as the right STG cluster is concerned, we successfully cross-decode valence across all conditions and groups, with the exception of E^‒^V (Figure 5d). In particular, while the coefficients relating brain activity to valence scores in congenitally deaf individuals can be used to cross-decode pleasantness in people with typical development (E^+^A, p-value < 0.001; E^+^M, p-value < 0.001; E^+^V, p-value = 0.001) and with congenital loss of sight (E^‒^A, p-value = 0.003), the opposite is true in the multisensory condition only (E^+^M, p-value = 0.045; E^‒^A, p-value = 0.062; E^+^A, p-value = 0.053; E^+^V, p-value = 0.054). Lastly, regarding the left STG cluster, the cross-decoding of valence is significant for all conditions and groups (Figure 5e).

## Discussion

In the present study, we explored how sensory experience and modality impact the neural representation of emotional instances, with the aim of uncovering whether the brain represents affective states through sensory-specific mechanisms or a more abstract coding. To achieve this, we employed a naturalistic stimulation approach encompassing either unimodal or multimodal conditions and collected moment-by-moment categorical and dimensional emotion ratings. We also recorded brain activity using fMRI from people with and without congenital sensory deprivation while presenting them with the same emotional stimuli.

Our results reveal that the ventromedial prefrontal cortex (vmPFC) represents emotion categories across modalities and regardless of sensory experience. In addition, we successfully decode the timecourse of emotional valence from the activity of the posterior portion of the superior temporal cortex (pSTS), even when participants lack visual or auditory inputs. Through multivariate pattern classification, we show that sensory experience, more than modality, can be decoded from regions of the emotion network, with the exception of vmPFC, which lacks any discernible information for decoding. Our data also reveal that early sensory areas represent the emotion model based on the stimulus modality. Specifically, in blind and typically-developed individuals exposed to the auditory movie, higher fitting values are observed in the early auditory cortex, whereas in the visual-only experiment, emotions are better fitted to the early visual areas of deaf and control participants. Lastly, higher-order occipital regions, like the fusiform gyrus, encode the emotion model similarly in both sighted and blind individuals when listening to the narrative.

Understanding whether mental faculties and their corresponding neural representation necessitate sensory experience for their development or whether they can independently form and evolve without such input is of paramount importance. Previous research has shown that the recruitment of cortical modules specific to object perception, action recognition, and spatial navigation, among others, can develop even in the case of congenital lack of sight (Pietrini et al., 2004; Ricciardi et al., 2009; Wolbers et al., 2011; Ratan Murty et al., 2020), supporting the idea of a more abstract coding of information within these regions. Although the study of sensory-deprived individuals provided crucial insights into the architecture of human cognition, no studies adopted the same approach to investigate whether emotions are represented in a supramodal manner rather than in a sensory-specific way.

We show through voxelwise encoding that vmPFC stores an abstract representation of emotion categories, as it is involved across sensory modalities and regardless of experience. To delve deeper into the role of this region, we examine with multivariate classification whether the pattern obtained from fitting the emotion model could effectively distinguish participants’ sensory experiences and the stimulus modality they are presented with. Multivariate results reinforce the notion of the supramodal nature of vmPFC, as information encoded in this area is not contaminated by sensory inputs. Previous research on typically-developed individuals only, already pointed to the recruitment of this region in processing emotional stimuli conveyed through the visual, auditory, tactile, and olfactory modalities (McCabe et al., 2008; Peelen et al., 2010; Chikazoe et al., 2014). In this context, examining sensory-deprived individuals becomes essential as it diminishes the likelihood that the recruitment of the same brain area across various modalities can be solely attributed to mental imagery (Pietrini et al., 2004). The vmPFC has been linked to the encoding of affective polarity and the regulation of emotional states (Winecoff et al., 2013; Hiser and Koenigs, 2018; Alexander et al., 2023). Also, lesions in this area alter emotional responses regardless of whether stimuli are presented through visual (Damasio et al., 1990) or auditory modalities (Roberts et al., 2004; Johnsen et al., 2009). Together with these findings, our data support the idea that vmPFC is crucial for generating affective meaning (Roy et al., 2012; Chang et al., 2021) and that such a meaning is represented in a modality-independent manner in both the sensory-deprived and the typically-developed brain. By obtaining categorical and valence ratings, we were also able to ascertain whether the abstract coding employed by vmPFC can be summarized by the dimension of pleasantness. Our findings indicate that the supramodal code of emotion in vmPFC is categorical rather than dimensional, as valence is mapped differently in typically-developed individuals listening to the movie as compared to all other groups and conditions. Consistent with prior findings highlighting the influence of imagery on the perceptual processing of emotionally charged stimuli (Diekhof et al., 2011), we posit that visual imagery may underlie the distinctive mapping of pleasantness in vmPFC of sighted. This proposition gains further support from the fact that congenitally blind individuals exhibit a valence representation comparable to that of sighted when exposed to the original movie or its muted version, along with evidence indicating their increased reliance on alternative forms of mental imagery (see Renzi et al., 2013 for a review).

Notably, a recent study investigating perceived voice emotions has revealed the coexistence of both categorical and dimensional accounts of affective experiences in the brain, where the activity of frontal regions represents emotional instances in categories, whereas temporal areas through dimensions (Giordano et al., 2021). Our findings not only reinforce but also expand the notion that the same emotional experience can be represented by distinct brain regions following either categorical or dimensional frameworks (Lettieri et al., 2019). Indeed, we show that the activity of pSTS tracks changes in valence regardless of the stimulus modality and in people with and without congenital sensory deprivation. This area is known for its involvement in processing emotions (Engell and Haxby, 2007; Basil et al., 2017), with its connectivity being predictive of affective recognition performance (Alaerts et al., 2014). Moreover, pSTS is crucial for the multisensory integration of emotional information from faces and voices (Hagan et al., 2009; Watson et al., 2014), as well as from body postures (Candidi et al., 2011; De Gelder et al., 2015). The activity of the superior temporal cortex is also modulated by the valence of vocal expressions (Frühholz and Grandjean, 2013) and facial movements (Narumoto et al., 2001; Jabbi et al., 2015). A prior study conducted on typically-developed individuals showed that the left pSTS maps emotion categories, irrespective of the sensory modality involved (Peelen et al., 2010). Our study confirms this finding, as we observed a convergence in left pSTS (x = -68, y = -14, z = -5) when examining the encoding results of the categorical model obtained from the audiovisual, visual-only, and auditory-only experiments conducted in people with typical development. However, we also find that the encoding of emotional instances in pSTS is influenced by sensory experience, as congenitally deaf participants do not show a mapping of emotion categories in the same region. In this regard, it is well known that the superior temporal cortex of deaf people undergoes a sizable reorganization so that deafferented auditory areas are recruited during the processing of visual stimuli (Vachon et al., 2013). Interestingly, the repurposing is more evident in the right hemisphere (Shibata et al., 2001; Finney et al., 2001; Sadato et al., 2005), and this might also account for our cross-decoding of valence from the left - but not the right - pSTS in congenitally deaf participants. These results collectively suggest that emotion category mapping in pSTS is contingent on sensory experience, as is the representation of valence in the right pSTS, whereas the left pSTS encodes pleasantness in a supramodal manner.

So far, our data indicate that the abstract coding of emotional instances is based on a categorical framework in vmPFC and a dimensional one in left pSTS. How does the brain encode and represent emotional experiences beyond vmPFC and pSTS?

Our multivariate classification results reveal that sensory experience more than the modality of emotion conveyance plays a role in how the brain organizes emotional information. Recent behavioral studies have demonstrated that blind individuals have distinct experiences related to affective touch (Radziun et al., 2023) and retain unique representations of bodily maps of emotions (Lettieri et al., 2023). Similarly, congenitally deaf individuals exhibit specific abilities in identifying musical happiness (Sharp et al., 2020). Notwithstanding these profound differences in the processing of affective stimuli, the precise mechanisms through which sensory deprivation shapes how emotional experiences are encoded in the brain remain to be fully elucidated. In our data, we show a significant mapping of emotion categories in higher-order occipital regions, such as the fusiform gyrus, in both congenitally blind and typically-developed individuals while listening to the movie. It is well known that the deafferented occipital cortex of blind people is recruited for perceptual (Collignon et al., 2011) as well as more complex tasks (Sadato et al., 1996; Bedny et al., 2011). Most of these studies discuss this evidence in the context of crossmodal reorganization (Merabet and Pascual-Leone, 2010). Here, instead, we show that the network of brain regions encoding emotion categories is similar between sensory-deprived and typically-developed individuals, even beyond areas representing affect in a supramodal manner. In line with this, we observe null results for the comparisons between people with and without sensory deprivation within modality. Hence, our data suggest that, as in the case of perceptual processes (Striem-Amit et al., 2015; Arcaro and Livingstone, 2017), the brain constructs a framework for the representation of emotional states, irrespectively from sensory experience. Sensory inputs during development, however, shape the functioning of this scaffold.

If the influence of sensory experience is notably pronounced, that of sensory modality is less conspicuous upon first examination. In fact, when considering how distinct brain regions represent our emotion model, it is not possible to discern the specific stimulus that is presented to typically-developed individuals. Nonetheless, we observe higher fitting values of the emotion model in the auditory cortices when participants are exposed to the audio version of the movie. Conversely, the video version is linked to higher fitting values in visual areas. This finding aligns with previous research, which has proposed the existence of what is known as emotion schemas within the visual cortex (Kragel et al., 2019). The concept of emotion schema suggests that visual features consistently associate with distinct emotions and contribute to our comprehensive emotional experiences. Our finding provides further support for the idea that emotions are not only processed in centralized supramodal emotional areas but also distributed across different sensory processing regions of the brain (Kragel et al., 2019; Čeko et al., 2022). The specificity of the relationship between sensory modality and elicited emotions is also testified by the differences we observe in the affective experiences reported by participants of our behavioral studies. Indeed, while amusement, love, joy, fear and contempt are the predominant emotions across our stimuli, fear is more challenging to experience solely through the auditory modality, while contempt is more easily elicited in the audio version of the movie. These results are consistent with previous studies demonstrating distinct emotion taxonomies associated with different sensory modalities to convey affective states (Cowen et al., 2019; Cowen and Keltner, 2020; Cordaro et al., 2020).

Of note, we believe it is relevant to acknowledge a possible limitation of our study, as we did not gather affective ratings from congenitally blind and deaf participants. Future investigations could address this by designing a novel methodology to capture real-time emotion reports in contexts of sensory deprivation.

In conclusion, we have shown that emotional experiences are represented in an extensive network encompassing sensory, prefrontal, and temporal regions. Within this network, emotion categories are encoded using an abstract code in the ventromedial prefrontal cortex and perceived valence is mapped in the left posterior superior temporal region independently from the stimulus modality and experience. Thus, sensory-specific and abstract representations of emotion coexist in the brain, so as categorical and dimensional accounts. Sensory experience more than modality impacts how the brain organizes emotional information outside supramodal regions, suggesting the existence of a scaffold for the representation of emotional states where sensory inputs during development shape its functioning.

